# Regulation of Age-Related Lipid Metabolism in Ovarian Cancer

**DOI:** 10.1101/2024.09.06.611709

**Authors:** Jihua Feng, Clay Douglas Rouse, Isabella Coogan, Olivia Byrd, Ethan Nguyen, Lila Taylor, Santiago Garcia, Hannah Lee, Andrew Berchuck, Susan K. Murphy, Zhiqing Huang

## Abstract

Although a lot of effort has been dedicated to ovarian cancer (OC) research, the mortality rate is still among the highest in female gynecologic malignancies. The effects of the aged tumor microenvironment are still being undermined despite age being the highest risk factor in ovarian cancer development and progression. In this study, we have conducted RNA sequencing and lipidomics analysis of gonadal adipose tissues from young and aged rat xenografts before and after ovarian cancer formation. We have found significantly higher tumor formation rates and volumes in aged OC xenograft rat models compared to their young counterparts (p<0.05), suggesting the aged adipose microenvironment (AME) is more susceptible to OC outgrowth. We have revealed significant shifts in the gene expression enrichment from groups of young vs. aged rats *before* tumor formation, groups of young vs. aged rats *when* the tumor formed, and groups of aged rats *before and after* tumor formation. We also observed shifts in the lipid components of the gonadal adipose tissues between young and aged rat xenografts when tumors were generated.

Additionally, we found that the aged AME was associated with age-related changes in the immune cell composition, especially inflammation-related cells. The top hits showing the most differences between aged and young adipose tissues were eight genes including S100a8, S100a9, Il1rl1, Lcn2, C3, Hba-a1, Fcna, and Pnpla3, 22 lipids including multiple isoforms of free fatty acids (FFA) and triglyceride (TG), as well as four immune cells including neutrophil, myeloid dendritic cell, T cell CD4+ (non-regulatory), and mast cell activation. The functional correlation among S100a8, S100a9, neutrophil, and FFA (18:3) was also determined. Furthermore, FFA (18:3), which was shown to be downregulated in aged xenograft rats, was capable of inhibiting OC cell proliferation. In conclusion, our study suggested that aging promoted OC proliferation through changes in genes/pathways, lipid metabolism, and immune cells. Targeting the aging adipose microenvironment, particularly lipid metabolism reprogramming, holds promise as a therapeutic strategy for OC, which warrants further investigation.

**Significance:** Aging microenvironment of OC may be regulated by S100a8 and S100a9 secreted by adipocytes, preadipocytes, or neutrophils through affecting the lipid metabolism, such as FFA (18:3).

## Introduction

Globally, ovarian cancer (OC) is the eighth most common cancer in women and is associated with the highest mortality among female gynecologic malignancies, 6.6% (324,398 new cases) of age-standardized incidence rate worldwide, and 4.0% (206,839 Deaths) of age-standardized mortality rate in females in 2022 [1]. This highlights the significant disease burden on women’s health and survival. Since OC tends to advance rapidly and exhibits no apparent signs or symptoms in its early stages, more than two-thirds of patients are diagnosed at an advanced stage [2]. OC is the most frequently diagnosed among women aged 55-64, with median age at diagnosis of 63 [3]. Two major factors that impact prognosis and survival are metastasis and recurrence, and these are known to be positively related to aging [4–6]. Furthermore, elderly patients have poorer prognoses, with unfavorable progression- free survival (PFS) and overall survival associated with increased age [3]. While treatment disparities in the elderly may contribute to this difference, the aged host may also be more susceptible to disease progression. The mechanisms that underlie the influence of age on OC development, progression, and recurrence are poorly defined.

OC seems to prefer spreading to the adipose tissue (AT) within the peritoneal cavity, specifically the omentum, in which the mechanisms related to the proliferation of metastatic OC cells in the intra-abdominal environment still lack understanding. The tumor microenvironment (TME) of OC comprises various cell types, including tumor and host stromal cells, blood vessels, and the extracellular matrix (ECM), whose roles in OC cell dissemination are complicated and uncertain even though some studies have suggested that the interaction between tumor and stromal cells may facilitate OC cell spread within the peritoneal cavity [7, 8].

There is increasing evidence showing that adipose tissues, the main TME components in OC, play an important role in promoting cancer initiation and progression [9]. OC cells closely interact with adipocytes in the omentum, leading to significant the phenotypic alterations in the adipocytes. These altered adipocytes, with different characteristics from those of primary adipocytes, are named cancer-associated adipocytes (CAAs). Cancer-associated adipocytes (CAA) are especially significant in cancer progression since they facilitate cancer cell growth, angiogenesis, anti- apoptotic effects, and migration directly or indirectly [10]. Furthermore, certain adipokines, such as leptin, interleukin (IL) 6, and IL-8, have been identified to contribute to the growth, spread, and chemo-resistance of various types of tumors, including OC. The current understanding is that CAAs are multifaceted and constantly evolving, having many roles in constructing a tumor-promoting TME [11]. As one of the biggest risk factors for cancer development, aging promotes cancer progression through a variety of mechanisms, including accumulated cellular damage, increased systemic inflammation, and attenuated adaptive immunity [12]. Adipose tissue displays species-conserved, temporal changes with ageing, including redistribution from peripheral to central depots, loss of thermogenic capacity, and expansion within the bone marrow [13]. Adipose tissue is among the earliest organs to respond to ageing, as determined by single-cell transcriptomic analyses, making it possible that it may subsequently drive whole-organism ageing [14]. Adipose tissue may therefore be a central driver for organismal ageing and age-associated diseases [13]. However, the extent to which these changes contribute to aging adipose tissue in aged OC remains unknown. Unfortunately, previous studies have been largely limited to the effects of aging in healthy donors or alternatively have focused on the impact of the TME in young mice [15], and the importance of age-adipose tissue in the aged TME remains unclear.

Therefore, our study focuses on the impacts of an aged adipocyte-rich TME by taking the adipose-enriched tissues surrounding OC xenograft rats on the cancer development and progression. The objectives of this study were to investigate how the gene regulation pathways and metabolic modification of aged versus young TME with a focus of adipose differ and how the aged TME contributes to a more aggressive OC phenotype. The tumor growth was compared between the young and aged rat groups. The gene expression/pathways, lipidomic, tumor immunity patterns, and the correlation of gene expression and lipid profiling were studied with the fat tissues surrounding the normal ovaries of young versus aged rats, the fat tissues adjacent to tumors pre- and post-tumor formation. A network of gene-lipid-immune cell infiltration in tumor-adjacent fat tissues in post-estropausal rats was established.

## Materials and Methods

### Cell lines

All ovarian cancer cell lines and Mouse preadipocyte 3T3-L1 cells line (Cat # CL- 173) were initially purchased from ATCC. OC cells were maintained in RPMI 1640 (ThermoFisher, Cat # A4192301) with 10% FBS (ThermoFisher, Cat# A5256801) and 1% Penicillin-Streptomycin (P/S, MilliporeSigma, Cat #p4333). 3T3-L1 cells were maintained in DMEM medium with high glucose (MilliporeSigma, Cat # D5796), 10% Bovine Calf Serum (BCS, MilliporeSigma, Cat # 12133C) and 1% P/S. Cells were incubated at 37°C in a humidified chamber with 5% CO_2_. All cell lines were genetically authenticated before using by the Duke University DNA Analysis Facility and confirmed to be free of mycoplasma by the Duke Cell Culture Facility.

### Senescence of 3T3-L1 Cells

Senescence of 3T3-L1 was induced by 2 μM carboplatin in the growth medium for 72 h according to literatures [16, 17]. The cells were washed with PBS and maintained in the growth medium for 24 h to recover. Then the cells were passed and incubated for 6 more days before use. The exposure algorithm was selected for the conditions that did not elicit robust cell death (<10–15%) while resulted in metabolic changes of cellular senescence.

### Beta-Gal Staining of 3T3-L1 Cells

The 3T3-L1 cells were fixed and stained with a senescence-associated β-galactosidase (SA-β-Gal) Staining Kit (Cell Signaling Technology, Inc, Cat#9860) according to the manufacturer’s instructions. Staining was performed in the cell culture petri-dish at 37°C overnight in an incubator without CO_2_. Images were taken under a microscope (EVOS M7000) at 10x10 magnification. The senescence levels were determined by the ratio of SA-β-Gal-positive cells with Image J by NIH. Average values from 5-6 images were taken for each well of 6-well plate. The whole procedure was repeated three times independently.

### Aged Conditioned Cell Culture and Cell Proliferation Assay

A2780 cells were cultured with the medium from young and aged 3T3-L1 cells using 96-well plate at 2000 cells/well. The carboplatin-unexposure 3T3-L1 cells (young conditioned medium, CM) were mixed with carboplatin-exposure 3T3-L1 cells (aged CM) at 1:2 ratio. The A2780 cell proliferation was analyzed 48h after cultured in CM using CellTiter One Solution Cell Proliferation Assay kit (MTS assay, Promega, Cat# G8461) according to the manufacturer’s instructions. The luminescence signal was recorded using the plate reader (BMG Labtech, POLARstar Omega). Readings were standardized to the cells cultured in young CM and reported as the percentage of viable cells. This test was repeated three times independently.

### FFA (18:3) Treatment

Human OC cell line A2780 and high grade serous epithelial OC cell lines, HEYA8 and CAOV2 were cultured with FFA (18:3), 9(Z),11(E),13(Z)-octadecatrienoic acid (aablocks, Cat# AA00DIYY) at the indicated dose ranges at 2000/well for A2780, 1000/well for HEYA8, and 2000/well for CAOV2 in 96-well plates. The cell proliferation was analyzed after 48 hours culture using CellTiter One Solution Cell Proliferation Assay kit (MTS assay, Promega, Cat# G8461) according to the manufacturer’s instructions. The luminescence signal was recorded using the plate reader (BMG Labtech, POLARstar Omega). Readings were standardized to the cells without FFA (18:3) treatment and were reported as the percentage of viable cells. This test was repeated three times independently.

### RT-qPCR

RNA isolation from 20 mg of adipose tissue was performed using TRI Reagent® (ThermoFisher. Cat# AM9738) according to the manufacturer’s instructions. RT-PCR was carried out using 300 ng of total RNA in a 20 µL of reaction using SuperScript IV One-Step RT-PCR kit according to the manufacturer’s protocol (ThermoFisher, Cat# 12594025) with Taqman probes specific to rat genes (Table S1 in supplementary data). Rat GAPDH was served as an endogenous control for RNA input. The PCR reaction was performed at 50°C 10 min for DNA synthesis followed by 95°C 1 min, 45 cycles of 95°C 10 seconds and 60°C 1 min. Relative RNA expression values were calculated using the CT values and normalized to GPADH CT values. This test was repeated three times independently with 2 replicates for each condition in each test.

### Rat Xenograft OC Model

The human ovarian adenocarcinoma cell line A2780 was used for rat xenograft model due to it has been successfully used for tumor generation in female athymic nude rats (RNU) rats by Charles River Laboratories. All animal experimental protocols were approved by the Duke Institutional Animal Care and Use Committee (IACUC). All methods are reported in accordance with ARRIVE guidelines. For “aged” (post-estropausal) group, 16 of 6-week-old RNU rats were raised in Duke University’s Division of Laboratory Animal Research (DLAR) animal facility until they reached the age of 12 months [18, 19]. During that span of time, the regular checks of body weight, health conditions, and behavior have been performed once a week. During this time span, 3 rats dead due to accident housing problem and left 13 rats in the aged study group. Sixteen RNU rats (6-weeks old) were purchased when aged rats reached to 12 months old. The surgery was performed to implant A2780 cells at 5x10^6^ per mouse embedded in 100 ul of Growth Factor Reduced Basement Membrane Matrix (Corning, Cat#354230) into aged rats and young rats. During this surgery, a piece of fat tissues termed as pre-tumor adipose (PTA) was removed and used for RNA-sequencing assays. After implantation, the rats were checked daily for tumor formation by visualization and palpate the abdomen. At 26 days after cancer cell implantation, all the aged and young rats were performed surgery to remove the tumors and a piece of fat tissues surrounding the tumor named as tumor surrounding adipose (TSA). Tumor length and width were measured, and the tumor volumes were calculated using the formular V = 0.5 × L × W^2^, where V is the tumor volume, L is the tumor length, and W is the tumor width.

### Transcriptome Sequencing (RNA-Seq)

RNA extraction with TRI Reagent® method (ThermoFisher) and RNA sequencing were performed by Admera Health with 20 mg of fat tissues, including 29 **PTA (**16 young and 13 aged rats) and 28 **TSA** (16 young rats and 12 aged rats) **tissues** collected as shown above in the section of Mouse Xenograft Model. The brief protocol was described in supplementary information.

### Quantitative Lipidomics Assay

The quantitative lipidomics analysis, which provided absolute quantification of up to 3000 lipid compounds, was performed at Metware Biotechnology Inc with the TSA tissues, including 13 adipose from aged rats and 18 adipose from young rats using ∼20 mg of the fat tissues per rat. The brief protocol was described in supplementary information.

### Statistical Analysis

The cell proliferation was evaluated using a student *t* test using GraphPad Prism 10 software (GraphPad Software, LLC). RT-qPCR for gene expression was analyzed and compared between groups using *Mann-Whitney* test. The tumor formation rate in rats was compared between the aged and young rat groups using *Fisher’s exact* test and the tumor volumes were analyzed using *Mann-Whitney* test. *P* values < 0.05 were considered statistically significant.

#### Transcriptome data analysis

All RNA-Seq analyses were performed in the Bioinforcloud platform (http://www.bioinforcloud.org.cn), using R package. Three groups of gene expression were compared, genes between aged (A) and young (Y) rats in PTA (PTA-A&Y), genes between aged (A) and young (Y) rats in primary tumor surrounding adipose (TSA-A&Y), and genes between PTA and TSA in aged rats (PTA&TSA-A) (**Figure 1A**). Differential gene expression (DGE) analysis was performed using DESeq2, with adjusted *P* valuesΣ0.05 were assigned as differentially expressed. The corrected *P*-values and |log2FC| were employed as thresholds for significant different-expression. The statistical enrichment of genes in the KEGG pathway and GO was analyzed using the Gene set enrichment analysis (GSEA) by KEGG data set and GO data set for rat species. DIOPT (Version 9.1) was used to convert rat genes into human genes (https://www.flyrnai.org/cgi-bin/DRSC_orthologs.pl). “Tumor IMmune Estimation Resource” (TIMER Version 2.0, http://timer.comp-genomics.org/, immune infiltration estimations by TIMER, CIBERSORT, quanTIseq, xCell, MCP-counter and EPIC algorithms) was used for systematical analysis of immune infiltrates across diverse cancer types to characterize the immune cell-type composition of multiple groups with varying gene expression profiles with PTA_A&Y, TSA-A&Y and PTA&TSA-A samples. The normalized quantified values were used as input to TIMER2.0. Differential infiltration levels of immune system cell types between the two comparison combinations were performed using the Limma in the Bioinforcloud platform and the immune cells with adjusted *P*Σ0.05 by Limma were assigned as differentially infiltration levels.

**Figure 1.**
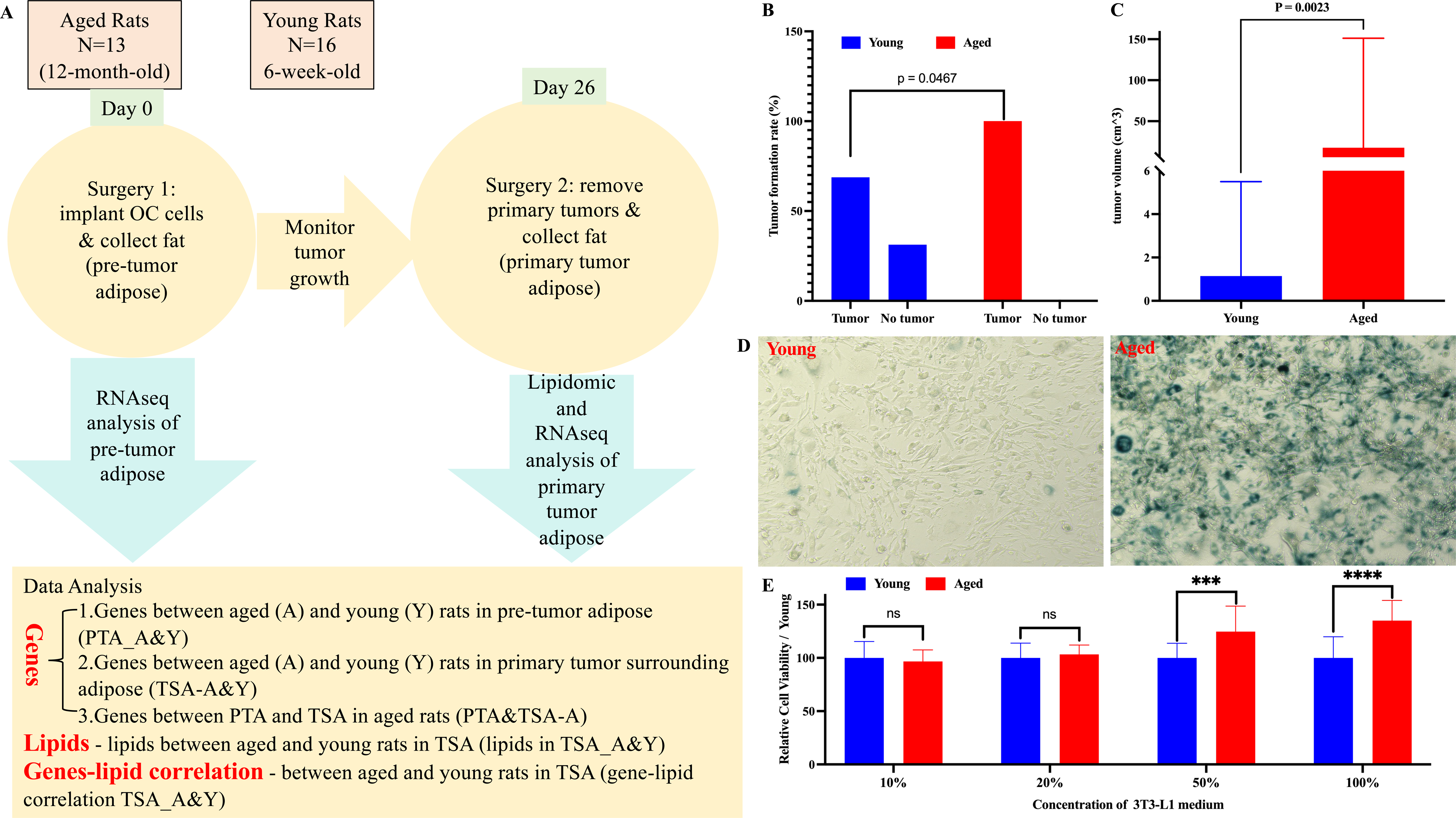
More tumor formation in aged TME. **A.** Workflow for xenograft rat model. **B.** Comparison of the tumor formation rate between aged and young rat groups. Thirteen out of 13 rats (100%) in aged group formed tumors by day 26 post OC cell implantation. Eleven out of 16 rats (68.75%) in young group formed tumors by day 26 post OC cell implantation. **C.** By day 26 post OC cell implantation, the average tumor volume from aged rat group was significantly bigger than that from young rat group, aged tumor median=17.64 cm^3^, young tumor median=1.14 cm^3^. **D.** Preadipocytes, 3T3-L1 were induced aging with carboplatin treatment at 2 μM for 72 hours. Blue color represents senescent cells. **E.** OC cell line, A2780 showed higher cell proliferation when exposed in an aging conditioned medium from aged 3T3-L1 cells. ^ns^ p>0.05, *** p<0.001, **** p<0.0001.

#### Statistical analysis of metabolome data

Orthogonal partial least squares discrimination analysis (OPLS-DA) and partial least square discriminant analysis (PLS-DA) were performed to obtain the VIP value of each metabolite from Lipidomic data. In the univariate analysis, the *P* value of each metabolite between the two groups was calculated using non-parametric test (Wilcoxon rank-sum test), and the fold change (FC value) of the metabolite between the two groups was calculated. The default criteria for differential metabolite screening were VIP≥1, *P*Σ0.05, and |log2FC|≥1. The RaMP-DB (integrating KEGG via HMDB, Reactome, WikiPathways) was utilized to examine the functions and metabolic pathways of the metabolites. The *P≣*0.05 was considered significantly enriched for the metabolic pathway. Metabolomic analyses were performed in the MetaboAnalyst 6.0 (https://www.metaboanalyst.ca/) for the lipids in TSA_A&Y.

#### Combined transcriptome and metabolome analysis

The correlations of gene-lipid, immune cells-gene, and immune cells-lipid in TSA_A&Y were carried out using Spearman’s correlation with Bioinforcloud platform.

#### OncoPredict for drug sensitivity analysis

The IC_50_ values for chemotherapeutic compounds were calculated for each TSA_A&Y sample according to the gene expression by RNA-Seq. OncoPredict R package was used to calculate the IC_50_ values [20] to predict drugs and biomarkers. The drug sensitivity was analyzed using unpaired *t*-tests and *p*<0.05 was considered as statistical significance.

## Results

### Aging is related to the tumor formation and growth

To explore the impact of aged host to tumor development, young (6-weeks) and aged (12-months) female athymic nude rats (RNU) were implanted intraperitoneally with A2780 cells at 5x10^6^ per rat and sacrificed primary tumors and surrounding fat tissues at day 26 post cancer cell implantation (**Figure 1A)**. **Figure 1B** showed 13 out of 13 (100%) aged rats formed the tumor while 11 out of 16 (68.75%) young rats formed the tumor (*p*=0.0467) by day 26 post cancer cell implanted. Furthermore, the tumor volumes in aged rats were significantly higher than that was in young rats with range of 0.75-223 cm^3^ and median of 17.64cm^3^ for tumor in aged rats and range of 0-21.5 cm^3^ and median of 1.14cm^3^ for tumor in young rats (*p*=0.0023, **Figure 1C**). To validate this finding, we performed OC cell growth study using aging conditioned medium (ACM) from aged preadipocytes 3T3-L1. While the aged 3T3-L1 cells showed the intensive beta-gal staining **(Figure 1D)** which suggested cell aging, A2780 showed higher cell proliferation when exposed to ACM from aged 3T3-L1 cells **(Figure 1E)**. Our data suggested that aged adipose microenvironment is more susceptible to OC outgrowth.

### Transcriptome analysis shows differential gene expression in PTA-A&Y, TSA- A&Y, and PTA&TSA-A

The transcriptome of gonadal adipose tissue from PTA-A&Y, TSA-A&Y, and PTA&TSA-A were analyzed by RNA sequencing (RNAseq). A total of 28,914 transcripts (**Table S2**) were subsequently analyzed using Principal Components Analysis (PCA) for gene expression. The score map (**Figure 2A**) showed the samples within each group (young or aged) were highly correlated and the distinguish between groups were relatively far, suggesting that there were differences in gene expression between the gonadal adipose tissue groups of PTA-A versus PTA-Y, TSA-A versus TSA-A-Y, and PTA-A versus TSA-A. When the threshold was set to |log2 FC|≥1 and adjusted p≤0.05, 2323 DEGs were identified between PTA-A and PTA-Y (DEGs), 1002 DEGs between TSA-A and TSA-Y, and 2612 DEGs between PTA-A and TSA- A (**Figure 2B**). The down- and up-regulated DEGs in aged group as well as no changed genes were also shown in **Figure 2C** for the comparisons of different groups. The heatmap in **Figure 2D** showed 25 up- and 8 down-regulated genes in aged rats for all three groups of comparisons of PTA tissues aged vs young, TSA tissues aged vs young, and aged PTA vs aged TSA. Among these genes, there were 7 (21.21%) adipokine-related genes including S100a8, S100a9, Mmp8, Lcn2, Pdpn, C3, and Pnpla3 (**Table S3**).

**Figure 2.**
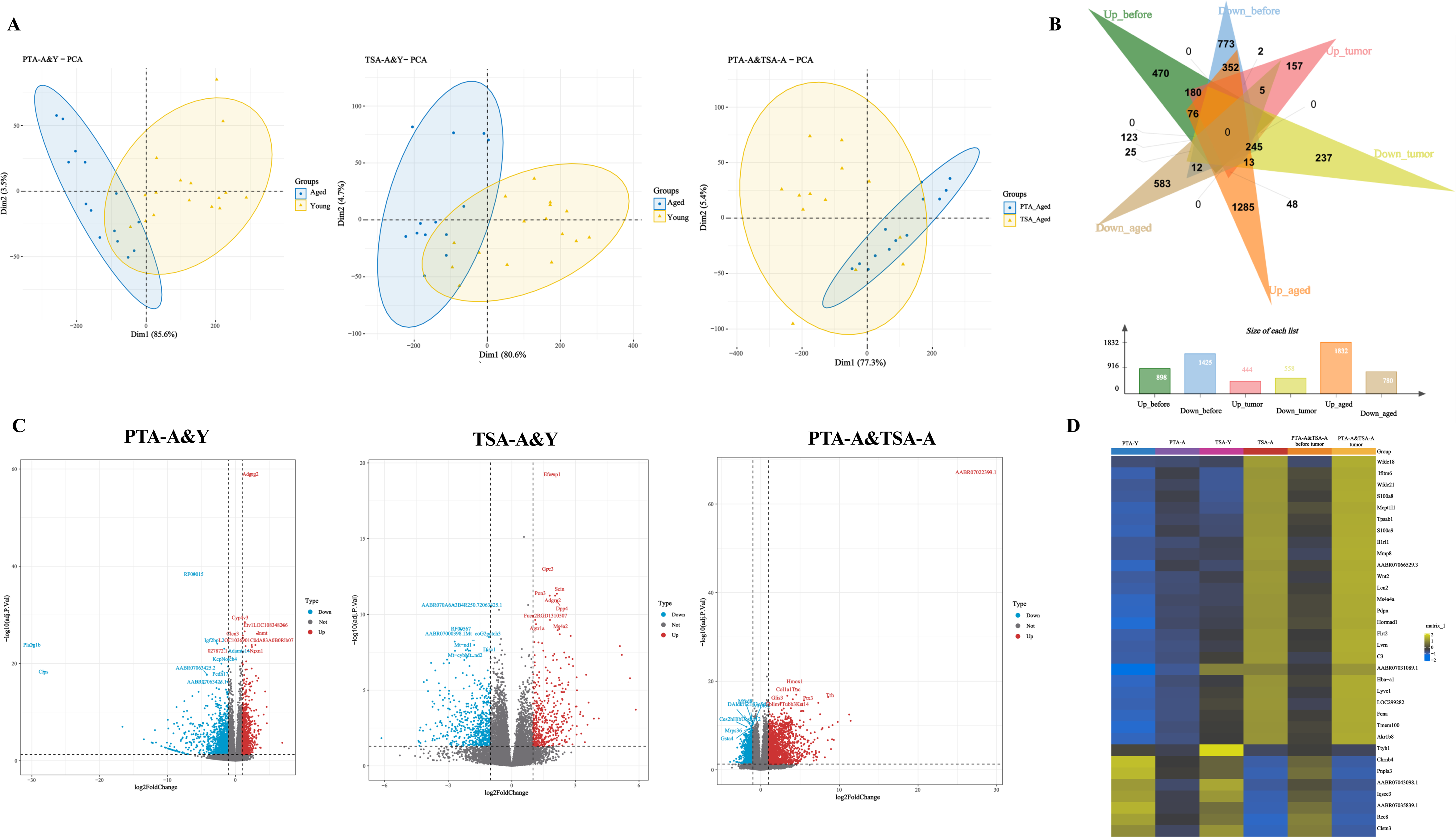
Transcriptome analysis showed differential gene expression in PTA- A&Y, TSA-A&Y and PTA&TSA-A. **A.** A principal component analysis (PCA) analysis was performed with 29 biological samples including 16 fat tissues from young rats and 13 from aged rats. The analysis showed the similarities of differential gene expressions (DEGs) within the aged (blue) or young (yellow) groups and the clear separations of gene expression between aged and young groups for all 3 panels of datasets, PTA-A&Y (left panel), TSA-A&Y (middle penal), and PTA&TSA-A (right penal). **B.** Venn plot of up-regulated and down-regulated DEGs in PTA-A&Y, TSA-A&Y, and PTA&TSA-A groups. Up-before: up-regulated genes in PTA-A&Y; down-before: down-regulated genes in PTA-A&Y; up-tumor: up-regulated genes in TSA-A&Y; down-tumor: down-regulated genes in TSA-A&Y; up-aged: up-regulated genes in PTA&TSA-A; down-aged: down-regulated genes in PTA&TSA-A. The bar graph showed the number of DEGs in each adipose group. **C.** Comparisons by Volcano plots of unique DEGs in PTA-A&Y, TSA-A&Y, and PTA&TSA-A groups.The x and y axes showed the log2 FC and –log10 (adjusted *P* value), respectively. Types of down, not, and up indicated down-regulated DEGs, no difference DEGs, and up-regulated DEGs in aged group. **D.** The heat map of DEGs in PTA-A&Y, TSA- A&Y, and PTA&TSA-A. The differential expression genes were shown on the right of the heat map.

Next, the KEGG pathway search and Go Analysis were carried out with the DEGs from 3 groups of comparison, PTA-A&Y, TSA-A&Y and PTA&TSA-A when the threshold was set to |NES|≥1 and *p*≤0.05. Twenty-six KEGG pathways, 45 biological process, 28 molecular functions, and 5 cellular components were differentially enriched from all three groups of comparison (**Figure 3**). These analyses revealed that several pathways/functions enrichment, such as Cytokine receptors, Cytokines and growth factors, Viral protein interaction with cytokine and cytokine receptor, IL-17 signaling pathway, and Glycerolipid metabolism. With the Go biological process analysis, we further identified the cytokine-mediated signaling pathway, fatty acid metabolic process, immune response, T cell activation, immunoglobulin mediated immune response, and negative regulation of T cell proliferation. This data suggested the involvement of cytokines and cytokine-related pathways, lipid metabolism, and immune cells in the OC development in aged animals.

**Figure 3.**
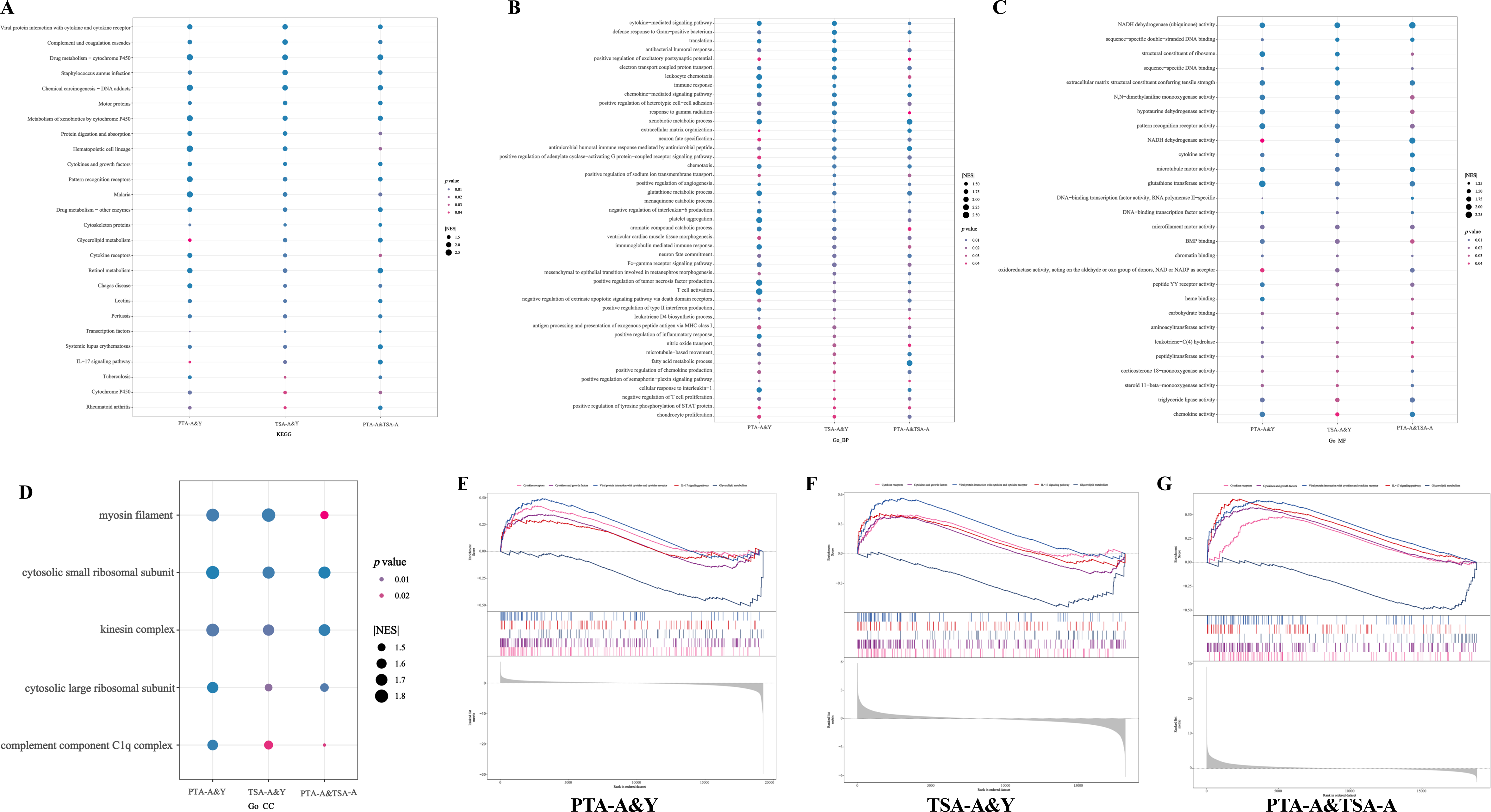
Enrichment of KEGG Pathway and Go functions in PTA-A&Y, TSA- A&Y, and PTA&TSA-A groups. A-D. Bubble charts of enrichment analysis by GSEA. Twenty-six KEGG pathways (**A**), 45 biological process (Go-BP, **B**), 28 molecular function (Go_MF, **C**), and 5 cellular components (Go-CC, **D**) were differentially enriched in comparisons of PTA-A&Y, TSA-A&Y, and PTA&TSA-A groups (|NES|*≣*1 & *p≣*0.05). NES: normalized enrichment score. Dot colors represented the levels of p values. The dot sizes represented the NES scores. **E-G.** Five of KEGG pathways were enriched in all 3 comparisons of PTA-A&Y, TSA- A&Y, and PTA&TSA-A groups: Cytokine receptors (pink lines), Cytokines and growth factors (purple lines), viral protein interaction with cytokine and cytokine receptor (blue lines), IL-17 signaling pathway (red lines), and Glycerolipid metabolism (black lines).

### Regulation of Immune Microenvironment of OC

TIMER was used to estimate the immune infiltration in all three comparison groups, PTA-A&Y, TSA-A&Y, and PTA-A&TSA-A. The various immune cell types were revealed significant differences in all three comparison groups, including 20 in PTA- A vs PTA-Y, 7 in TSA-A vs TSA-Y, and 30 in PTA-A vs TSA-A (**Table S4**). The higher infiltration levels were shown for neutrophil, T cell CD4+ (non-regulatory), and myeloid dendritic cell in aged rats compared with young rats in PTA and TSA, as well in TSA-A compared with PTA-A. The infiltration levels of mast cell activation were significant different between the aged groups with higher infiltration levels in aged rats compared with young rats before tumor generation (PTA-A vs PTA-Y), and aged OC xenografts compared with young OC xenografts when primary tumor formed (TSA-A vs TSA-Y). The lower infiltration levels of mast cell activation in aged rats after OC generation (TSA-A) were shown when compared with them before tumor formation (PTA-A).

### Lipid metabolic involvement in aged tumor generation

The metabolome OPLS-DA analysis from rat fat tissues at primary tumor formation were showed in **Figure 4A**. A total of 28 TSA samples including 13 aged and 15 young were clearly distinguished by the orthogonal T score 35.1 % and T score 5.1 % of the total variability (**Figure 4A**), respectively. T scores for the repeated samples within each group (young or aged) were closely related, and they were separated clearly between the groups, indicating the reliability of the metabolome data. A total of 905 metabolites in the metabolome were identified (**Table S5**), including 545 positive ions and 360 negative ions. According to RaMP functional annotation, 70 pathways were enriched, of which the largest number of metabolites was 24 (34.29%). The top 25 hits of functional pathways were shown in **Figure 4B**, which included the top 10 relevant pathways, such as G alpha (q) signaling events; Synthesis, secretion, and inactivation of Glucagon-like Peptide-1 (GLP-1); Incretin synthesis, secretion, and inactivation; Free fatty acid receptors; Linoleic acid oxylipin metabolism; Peptide hormone metabolism; Class A/1 (Rhodopsin-like receptors); GPCR downstream signaling; Octadecanoid formation from linoleic acid; and Omega-3/omega-6 fatty acid synthesis. With the threshold of VIP≥1, |log2 FC|≥1, and *P*<0.05, 37 differential expression of metabolites (DEMs) between young and aged groups were selected, including 17 up-regulated lipids and 20 down-regulated lipids in the aged rat when compared with the young rats at the stage of primary tumor formation (**Figure 4C, Table S6**). The top 10 significant DEMs were: CoenzymeQ8, CoenzymeQ9, Taurocholic acid, Triglycerides (TG) (18:1_20:1_24:1), TG (19:0_18:1_20:1), TG (18:1_20:1_22:5), PE (19:0_18:2), TG (24:1_18:2_18:2), TG (18:1_18:1_26:1) and TG (24:0_18:1_20:1). **Figure 4 D-F** showed the top 3 DEMs, including 13 Glycerolipids (GL), 10 Glycerophospholipids (GP), and 9 Fatty Acyls (FA).

**Figure 4.**
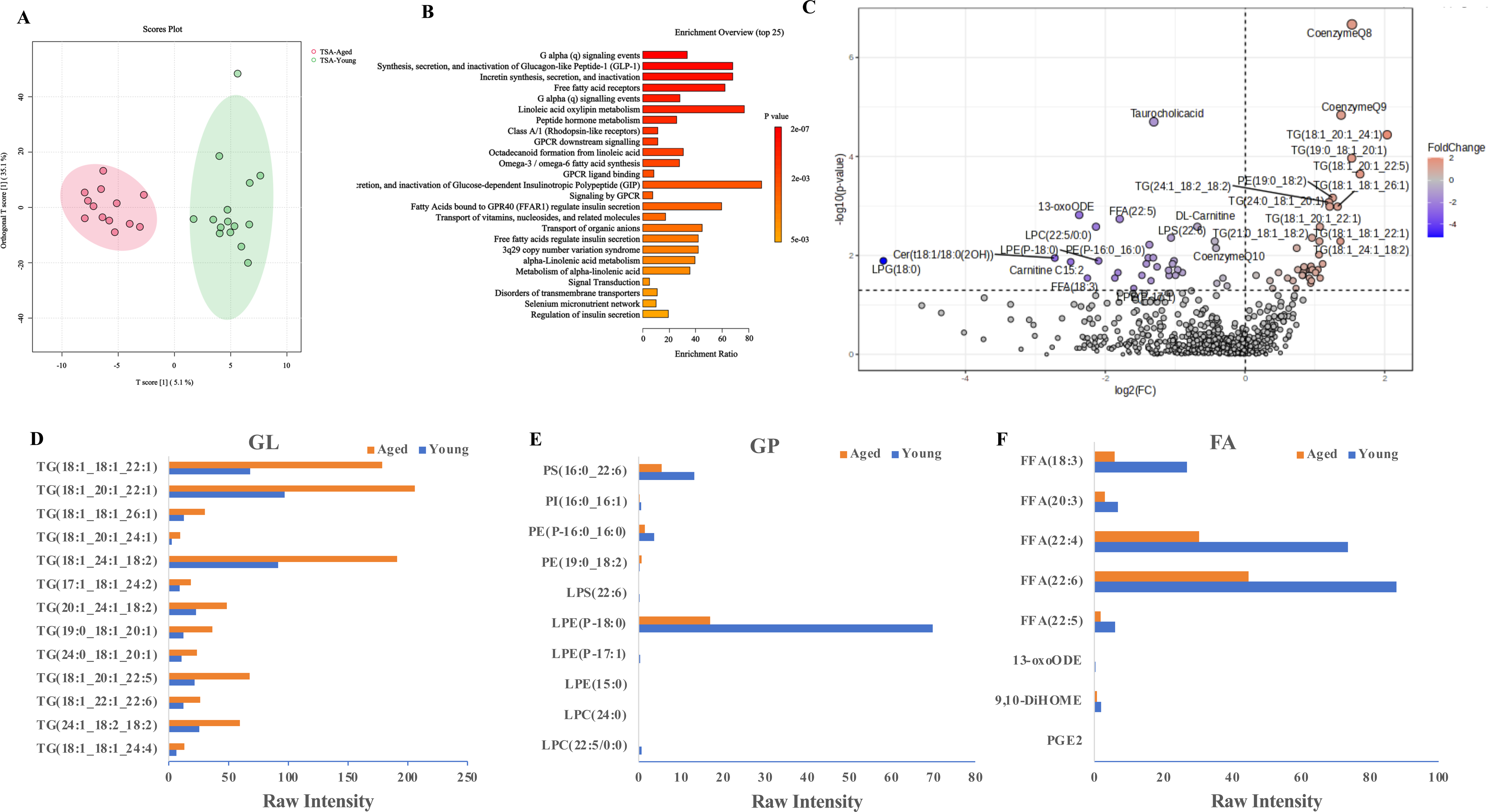
Differential expression of metabolites (DEMs) in TSA-A&Y adipose. **A.** Orthogonal Partial Least Square Discriminant Analysis (OPLS-DA) was carried out with 28 biological samples including 13 adipose from aged rats and 15 adipose from young rats at primary tumor formation. Green plot showed the lipid components in young rats and pink showed the lipids in aged rats. The lipid samples within each group were closely related, while the separation of lipids between the groups was far, indicating the reliability of the metabolome data from the aged rat adipose groups. **B.** The top 25 pathways of RaMP functional annotation. **C.** Volcano plots showed differential expression of metabolites (DEMs) between TSA-A versus TSA-Y. The x axes showed the log2 FC and y axes showed -log10 (*P* value), respectively. **D-F.** The bar graphs showed scaled intensity values of altered Glycerolipids (GL, **D**), Glycerophospholipids (GP, **E**) and Fatty Acyls (FA, **F**) between TSA-A (orange bar, Aged) versus TSA-Y (blue bars, Young).

### Transcriptome-metabolome newtworks

We next analyzed the data to explore the correlation between gene expression from RNA-seq assay and lipid metabolites assay from Lipidomic with the tumor surrounding adipose (TSA) tissues when primary tumor generated. With threshold of Correlation r≥0.5, |log2 FC| ≥1 for gene expressions, and *P*<0.05, 207 DEGs and 67 DEMs were identified (**Figure 5A**). Among these, punicic acid (9E,11Z,13E)- 9,11,13-Octadecatrienoic acid (FFA (18:3)) and PS (16:0/22:6 (4Z,7Z,10Z,13Z,16Z,19Z)) a phosphatidylserine showed the best explanatory variable in the fatty acid metabolic process and Glycerolipid metabolism (correlated to Pnpla3 gene). LysoPE (0:0/22:5(4Z,7Z,10Z,13Z,16Z)) (LPE (0:0/22:5)), a lysophosphatidylethanolamine or a lysophospholipid was the best explanatory variable in the Cytokine receptors (correlated to gene). FFA and TG were the best explanatory variables in the IL-17 signaling pathway shown correlated to genes S100a8 and S100a9. FFA (18:3) was correlated to genes of Pnpla3, S100a8, and C3. A more detailed description of the variables was available in **Table S7** in Supplementary Information.

**Figure 5.**
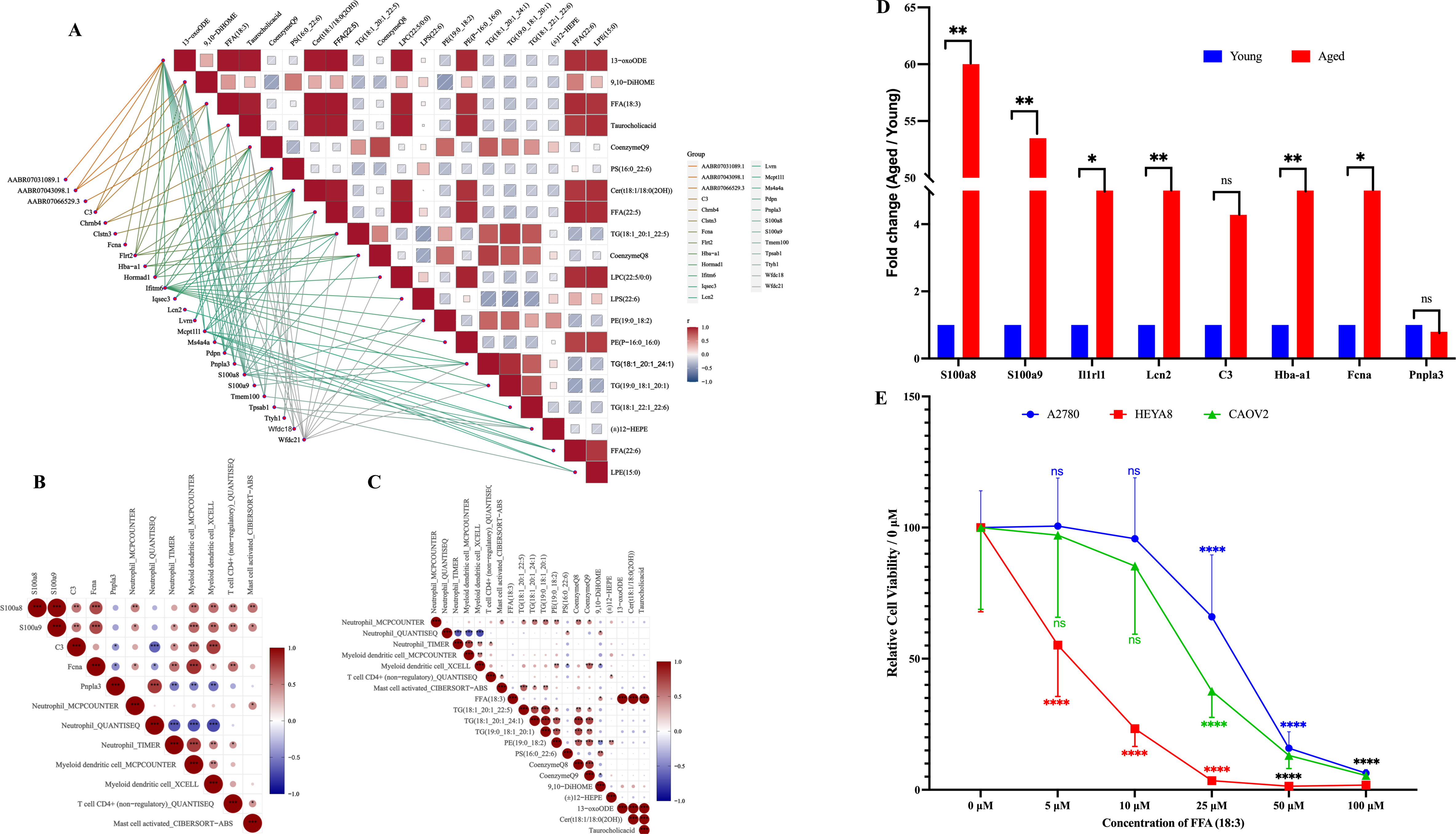
Transcriptome-Metabolome Correlative Regulation. **A.** The gene expression profiling from RNA-seq analysis and lipid profiling from lipidomic analysis were analyzed for their correlation in groups of TSA-A&Y, the adipose from primary tumor surrounding tissues from aged and young rats. With the thresholds of correlation ≥ 0.5, absolute log2 FC ≥ 1, VIP ≥ 1, *p* <0.05, we found 25 genes and 20 lipids that are closely related. The darker the colors showed the higher the correlation coefficient between gene expression and lipids. **B-C**. The correlation between immune infiltration and gene expression from RNA-seq and lipid metabolites from Lipidomic in groups of TSA-A&Y. **B.** The correlation between 4 immune cells (Neutrophil, Myeloid dendritic cell, T cell CD4+ (non-regulatory) and Mast cell activated) and 5 genes (S100a8, S100a9, C3, Fcna, Pnpla3). **C.** The correlation between 4 immune cells and 13 lipids. **D.** Gene expression was validated using Rt-PCR. Eight genes showed differential expression in groups of PTA-A&Y, TSA-A&Y and PTA&TSA-A and were involved in lipid regulation were validated using Rt-PCR with the gene specific probes for Taqman assay and 6 adipose from each of aged rats and young rats. As shown with RNA-seq analysis, the genes of S100a8, S100a9, C3, Il1rl1, Lon2, Hba-a1 and Fcna showed upregulation in aged rats and gene Pnpla3 showed downregulation in aged rats. **E.** OC cell line, A2780, HEYA8, and CAOV2 showed lower proliferation when exposed in FFA (18:3) containing medium in a dose dependent manner. ^ns^ p>0.05, **** p<0.0001.

We also explored the correlation between immune infiltration and gene expression from RNA-seq and lipid metabolites from Lipidomic assay from the tumor surrounding adipose (TSA) tissues when primary tumor generated as shown in **Figure 5B** for gene-immune cell infiltration and **Figure 5C** for Lipid- immune cell infiltration. With threshold of Correlation r≥0.5 and *P*< 0.05 for gene versus immune infiltration and lipid versus immune infiltration, we found that the infiltration levels of neutrophil were related to the expressions of genes S100a8, S100a9, C3, Fcna and Pnpla3, and lipids 13-oxoODE, Cer(t18:1/18:0(2OH)), CoenzymeQ8, CoenzymeQ9, PE(19:0_18:2), Taurocholicacid, TG(18:1_20:1_22:5), TG(18:1_20:1_24:1), TG(19:0_18:1_20:1), 9,10-DiHOME, and FFA(18:3). The infiltration levels of T cell CD4+ (non-regulatory) were related to genes S100a8 and S100a9, and lipids CoenzymeQ8, CoenzymeQ9, Taurocholicacid and TG (18:1_20:1_24:1). The infiltration levels of myeloid dendritic cell were related to genes S100a8, S100a9, C3, Fcna and Pnpla3, and lipids 9,10-DiHOME, (±)12-HEPE, 13-oxoODE, Cer (t18:1/18:0(2OH)), FFA (18:3), CoenzymeQ8, CoenzymeQ9, PE (19:0_18:2), Taurocholicacid, TG (18:1_20:1_22:5) and PS (16:0_22:6). The infiltration levels of mast cell activation were related to genes S100a8 and S100a9, and lipids CoenzymeQ8, CoenzymeQ9, TG (18:1_20:1_22:5), TG (18:1_20:1_24:1) and TG (19:0_18:1_20:1). Additionally, FFA (18:3) was correlated to Neutrophil, and Myeloid dendritic cell. A more detailed description of the variables was available in **Table S8** in Supplementary Information.

To validate the gene expressions from the RNA-seq analysis, we performed RT-qPCR for the specific genes. These included all 8 genes shown significant involvement in lipid regulations, such as Glycerolipid metabolism by Pnpla3 gene, Cytokine receptors by Il1rl1 gene, IL-17 signaling pathway by genes S100a8, S100a9, and Lcn2. With consistent to the RNAseq data, the genes S100a8, S100a9, C3, Il1rl1, Lon2, Hba-a1, and Fcna showed up-regulation while Pnpla3 gene down-regulation in fat tissues from aged rats compared with the fats from young rats (**Figure 5D**).

In order to further determine the functional contribution of lipids in OC cell growth, we test FFA (18:3) in cell proliferation of OC cell. Three OC cell lines were tested, including A2780, HEYA8, and CAOV2, with FFA (18:3) added into the cell culture medium at the concentration indicated in the **Figure E**. We have addressed a down- regulation of FFA (18:3) in aged rats from the lipidomic analysis which suggested a negative correlation with the more aggressive phenotype observed in aged animals (**Figure 1B, 1C**). With an agreement with the *in vivo* data, addition of FFA (18:3) resulted in a less OC cell proliferation (**Figure 5E**) in all 3 cell lines tested.

### Chemo-responses were correlated to aged TME

To identify more specific drug candidates for aged OC treatment, we used OncoPredict R package, a ridge regression-based method to predict the potential drug responses. With threshold *P*<0.05 in *t*-test, 27 drugs were predicted the potential differential responses between aged and young rats with OC. Among these, the IC_50_ values of 26 reagents were significantly lower in the aged group than that in the young group, suggesting the more chemo-sensitivity and their potential more effective treatment for the aged OC patients (**Table 1**). Whereas the sensitivity of Cediranib (**Table 1**, red lettered) showed an opposite sensitivity with higher IC_50_ values in the aged group suggesting the more resistant feature in aged patients.

## Discussion

This study aimed to explore the functional contribution of aged tumor microenvironment with a focus on adipose tissues in cancer development. We performed the study using 16 of 6-weeks old rats as young study group and 13 of 12- months old rats as the aged study group. The tumor generation was compared between two groups at day 26 post cancer cell implantation. As expected, the tumor formation in aged animals were more than young animals with more tumor formation rates (100% for aged rats vs 68.75% for young rats, *p*=0.0467) and larger tumor volumes (*p*<0.0023). The fat tissues around the ovaries before tumor formation and tumor surrounding fat tissues when primary tumor formed were dissected and analyzed using RNA sequencing and lipidomic to search for the genes and lipids contributing to tumor development in rat model. We have found that genes S100a8, S100a9, C3, Il1rl1, Lon2, Hba-a1, and Fcna were significantly higher expressed in aged animals, and Pnpla3 was lower in aged animals. Importantly, we have identified expression of specific genes that are involved in lipid regulation in aged animals, such as FFA (18:3) with genes S100a8, C3, and Pnpla3, LPE with gene Il1rl1, FFA and TG with genes S100a8 and S100a9. We have also identified the increased infiltration levels of immune cell, such as neutrophil, T cell CD4+ (non-regulatory), and myeloid dendritic cell, in the aged OC models as compared with the young animals. The correlation between immune cell infiltration and gene expression have been determined as well, such as neutrophil were related to gene expressions of S100a8, S100a9, C3, Fcna and Pnpla3, T cell CD4+ (non-regulatory) were related to genes of S100a8 and S100a9, and myeloid dendritic cells were related to genes S100a8, S100a9, C3, Fcna, and Pnpla3. Finally, we used the gene expression profiling from RNAseq assay and identified 26 chemotherapeutic drugs are more specific for aged OC treatment.

Ovarian cancer is a cancer of postmenopausal women, and it is rare in women below the age of 40 years. This study used 12-month-old rats to mimic postmenopausal women because studies have shown that rats typically experience irregular estrous cycles (termed estropause) around the age of 9-12 months due to the disruptions of hormones and a resulting reproductive decline [18]. Athymic nude rat xenograft model provides several advantages in aging and metastasis studies. Compared to the typical athymic nude mouse xenograft model, the rat model is better suited to our objectives with advantages including larger abdominal fat pads (a common site of OC metastasis), more robust chemotherapy and surgical tolerance, longer lifespan, ability to harvest a larger volume of blood/tissue, and pharmacokinetics that are more closer to humans [21]. More importantly, the metabolic physiology of the rat is more closely aligned with that in humans than is in mouse [22], making the research findings more translatable to humans. The larger size of rats allows for more sophisticated surgical manipulations and allows for collection of more abundant tissues and samples, reducing the number of animals required. In addition, Athymic nude rats were used in this study due to the wide acceptance as a model organism and are able to grow a large number of tumor cell lines of human and rodent origin for several months [23]. Therefore, we used 6-week-old female athymic nude rats , which have shown to be sexually mature [24], as the young study group and 12-month-old rats, which have shown post-estropause, as the aged study group.

The tumor microenvironment (TME) is a heterogeneous ecosystem composed of infiltrating immune cells, mesenchymal support cells, and matrix components contributing to tumor progression. Adipose tissue, a primary cellular component in TME, has not been studied deeply in the progression of solid cancers, especially in OC; thus, it has become a focus of our study. We first found more tumor formation and larger tumor volumes in aged animals than in young animals. This finding was also supported by an *in vitro* study using cell culture model, which showed that the senescent preadipocytes (3T3-L1) promoted more proliferation of OC cells. Our data suggested the aged tissue microenvironment, primarily adipose, provides favorable “soil” to the cancer development and growth. Studies have shown that the infiltration of inflammatory cells and the enlargement of Lipid droplets (LDs) in adipocytes are more sever in aged adipose tissues [25]. Compared to mature adipocytes, aged adipocytes exhibit an increase in senescence and a significant enhancement in the expression of pro-inflammatory cytokines and ECM components [26]. The progenitor cells (also called adipose tissue stem cells or stromal progenitor cells) in adipose tissues are among the most responsive to ageing and ageing related environment [13]. Adipose tissue-derived mesenchymal stem cells from older donors (human and mouse) exhibit senescence properties and impaired regenerative potential [27, 28]. In addition, although increasing adipocyte size allows for increased lipid storage with ageing, the larger adipocytes are more insulin resistant compared to smaller adipocytes, leading to enhanced rates of basal lipolysis [29]. Inefficient storage of lipids and increased basal lipolysis in hypertrophied adipocytes normally results in exposure of vicinal non-adipose cells to increased levels of fatty acids. Excess fatty acids have shown the negative impact on systemic energy homeostasis. This imbalance in energy homeostasis may favor tumor growth in older humans.

Increasing studies also suggested the aged host is more susceptible to cancer metastasis [4] , which on the other hand, support our findings that more aggressive cancer has shown in aged animals. Our studies provided strong evidence of the molecular mechanisms behind those cellular changes.

Lipid metabolism is a fundamental source of cellular energy, but its metabolism through mitochondrial beta oxidation results in enhanced redox stress, and impaired beta oxidation results in accumulation of toxic by-products that can result in cell senescence and aged TME. Due to the lack of studies on how metabolic reprogramming with aging has emerged as a critical feature on cancer development, we used high-throughput lipid metabolomics to characterize lipid metabolism in aged ovarian cancer. We have found the important lipid components, such as TG, PE, FFA, and coQ8/coQ9 were all different in aged animals compared with the young animals suggesting their contribution to the more aggressiveness of cancer phenotypes in aged animals. Interestingly, unlike other lipids that are most able to promote tumor development, we found that FFA (18:3), which showed downregulated in aged rats (Figure 4F), inhibits OC cells proliferation (Figure 5E). FFA (18:3), (9E,11Z,13E)- 9,11,13-Octadecatrienoic acid, is a punicic acid, also known as 9t,11C,13t-CLN or punicate, belongs to the class of organic compounds known as lineolic acids and derivatives (https://hmdb.ca/metabolites/HMDB0030963). FFA (18:3) has been reported to have anticancer properties on various cancer cells, such as breast cancer cells (MCF-7), lung cancer cells (A549), colorectal cancer (DLD1), stomach cancer cells (MKN-7), and Liver cancer (HepG2) [31]. FFA (18:3) can induce ferroptosis in diverse cancer cells as a single agent and does so through triggering ferroptosis by a mechanism distinct from canonical ferroptosis inducers [30]. Chou *et al.,* reported that the lower adipogenic effect of FFA (18:3) in 3T3-L1 cells mediated by eliciting apoptosis of preadipocytes [31]. The specific cytotoxicity of FFA (18:3) for preand post confluent preadipocytes and the evoked apoptosis might contribute to the downregulation of adipocyte differentiation [31]. We found that aged 3T3-L1 promoted the proliferation of OC cells compared with non-senescent 3T3-L1 (Figure 1E). Whether FFA (18:3) downregulation in the aged TME is due to reduced synthesis in senescent 3T3-L1 cells requires further investigation. Taken together, FFA (18:3) has significant potential as the chemo-preventive and therapeutic agents for cancer, which warrants further research to assess their mechanism of action and potential applications clinically.

In addition to exploring lipid metabolism in aged OC, we also explored the gene transcriptome of the TME of aged OC. We have identified the significant differential gene expressions (DEGs) in rat adipose tissues in 3 categories, without tumors (PTA- A&Y) and with tumors (TSA-A&Y) between aged rats versus young rats, and aged rats without tumors and with tumors (PTA-A vs TSA-A). Previous single-cell transcriptomics and RNA-sequencing studies have shown that differential gene expression (DEGs) arises in mid-life in gonadal adipose tissue (GAT) organs before other organs and persists with advanced age, exhibiting firm aging profiles [14]. This suggests that the adipose gene’s change is an early event in life and sensitive to aging. Aging-related genes in adipose tissue can facilitate cancer cell proliferation, and cancer cells, in turn, can deteriorate the TME, forming a vicious cycle. In this study, 25 up- and 8 down-regulated genes were differentially expressed in all three category comparisons PTA tissues A vs Y, TSA tissues A vs Y, and PTA-A vs TSA-A, indicating their functional contributions to adipose aging and tumor development in aged individuals. Correlation analysis further connected the DEGs and DMGs and emphasized the importance of DEGs in lipid metabolisms. Setting the stringent thresholds for correlation r value (Cor) and p-value, we obtained 8 candidate genes (S100a8, S100a9, IL1RL1, Lcn2, C3, Hba-a1, Fcna and Pnpla3) and 22 candidate lipids in the adipose of TME surrounding OC tumors. Adipocytes can secrete >600 metabolites, hormones and cytokines, collectively known as adipokines [32]. Our candidate genes are most adipokine-related genes except for Il1rl1 (Interleukin 1 Receptor Like 1) and Hba-a1, whose functions are related to proinflammation and abnormal aerobic respiration, respectively. Calprotectin, S100a8 and S100a9, are calcium- and zinc-binding proteins belong to the S100 family. They play a prominent role in the regulation of inflammatory processes and immune response in diseases including cancer. They are abundant in cytosol of neutrophils and can induce neutrophil chemotaxis and adhesion. The S100a8, S100a9, and S100a8/a9 heterodimer have been reported as factors involved in cancer development, progression, and metastasis by interfering with tumor metabolism and the microenvironment [33]. The high expression of S100a8 and S100a9 has been also found in ovarian cancer [34, 35]. Our results further suggested S100a8 and S100a9 are higher expressed in aged TME of ovarian cancer and could be associated with more aggressive cancer phenotype from the aged animals. Previous studies suggested that S100a8/a9 heterodimer can bind to polyunsaturated fatty acids such as oleic, linoleic, and arachidonic acid in a calcium-dependent manner [36]. Supportively, our data has shown that the expression of S100a8/a9 in TME was correlated with TG and various fatty acids, such as FFA (12:0), FFA (15:0), and FFA (18:3), in aged animals. Future work will be focused on how cancer cells are influenced by the TME’s S100a8/a9 regulated lipids. The involvement of TME’s S100a8/a9 in cancer cell metabolism via mitochondrial functions of cancer cells will be explored. Moreover, studies have suggested that neutrophils can induce the reactivation of dormant tumor cells (cancer stemlike cells) via the release of S100A8/A9 proteins and modified lipids [37], which further emphasized the importance of TME’s S100A8/A9 proteins on cancer development and recurrence through cancer stem cells. Unraveling molecular mechnisms mediated by S100A8/A9 would provide exciting opportunities to study and design novel cancer therapeutics. Other adipokine-related genes, such as Lcn2, C3, Fcna, and Pnpla3, were also discovered in our study for their association with the lipids in the aged OC model. These genes are worthy of further study for their functions in OC genesis.

Chronic inflammation is an important feature of aging. It can accelerate disease progression in many types of cancer and often exacerbate during conventional cancer treatments. Our data revealed that age-related changes in the immune cell composition, especially inflammation-related cells, in adipose tissue may contribute to cancer proliferation efficiency in aged animals. Furthermore, the relationships of gene expression, lipid components, and immune cell composition have also been demonstrated suggesting their functional connections in aged cancer. The lipid changes resulted by aging-related gene expression (such as S100a8/a9) could initiate immune cell infiltration, which shift towards a pro-inflammatory phenotype and promote age-related adipose tissue inflammation in cancer [38, 39]. This is supported by the evidence of an inflammatory pattern is present in obese and ageing adipose tissue with the abundant accumulation of M1 macrophages, neutrophils, mast cells, B2 cells, CD8+ T cells, and Th1 cells [40]. Indeed, we have observed that immune infiltration level of neutrophil, myeloid dendritic cell and T cell CD4+ (non-regulatory) and mast cell activation was higher in aged rats before and after OC cells xenotransplantation compared to young rat. Specifically, we found that higher expression of S100a8 and S100a9 were related to increased infiltration levels of neutrophil, myeloid dendritic cell, T cell CD4+ (non-regulatory), and mast cell activation. The lipid FFA (18:3) was correlated to the infiltration of Neutrophil, and Myeloid dendritic cell. Studies have shown neutrophils facilitate ovarian cancer premetastatic niche formation in the omentum [41]. While other study suggested that neutrophil extracellular trap formation correlates with favorable overall survival in high grade ovarian cancer [42]. Future studies on tumor-infiltrating neutrophils and its regulation in aged cancer should be performed.

In preclinical studies, the contribution of the aged microenvironment to therapy response is largely ignored, with most studies designed using 8-week-old mice rather than older mice that reflect an age appropriate to the disease being modelled. This may explain, in part, the failure of many preclinical therapies upon their translation to the clinic. We identified several chemo-drug candidates that potentially more effective for aged OC treatment. Future tests will be performed for their age-related anti-tumor effects both *in vitro* and in *vivo*.

However, there are several limitations that necessitate further investigation. Firstly, the major disadvantage of using athymic nude rats as a model system is that their lack of a functional immune cells necessarily affects the TME in the omental fat.

Secondly, this study used 12-month-old rats as the aged animals because, around the age of 9-12 months, rats typically experience irregular estrous cycles (termed andropause) due to hormone disruptions and a resulting reproductive decline [18].

Future studies using even older animals, such as 2-years-old, are critical to mimic human OC patients who are primarily affected in their 60s. Thirdly, as the preliminary step, we have reported a network of genes-lipids-immune cells in aged OC rats.

Further mechanistic studies will be performed using both *in vivo* and *in vitro* models. Additionally, the anti-tumor effects of chemotherapy candidates for aged OC treatment will be explored in the future.

In summary, this study is the first to depict the cancerous aging adipocyte-rich microenvironment landscape, including its gene characteristics, tumor immunity patterns, and lipid metabolism. We have demonstrated that the aged host is more susceptible to OC proliferation in both *in vitro* and *in vivo* aged models. Using the high-throughput strategies for gene expression and lipid profiling, we have identified the functional network of gene-lipid metabolism-immune cells in aged ovarian cancer.

S100a8 and S100a9 secreted by adipocytes or preadipocytes or neutrophils showed as the key genes affecting lipid metabolism, such as FFA (18:3), and in turn the cancer development in elders. This study is of great importance because it provides enriched genetic and metabolic information for the aging global population, and the number of older women diagnosed with OC is expected to rise. The strong evidence gained from this study suggests that interventions to maintain a functional microenvironment may restore age-related adipose tissue dysfunction, such as by disturbing TME through gene expression in lipid profiling, and could provide more effective treatment opportunities in older patients with ovarian cancer.

## Disclosure of Potential Conflicts of Interest

All authors declare that they have no known competing financial interests or personal relationships that could have appeared to influence the work reported in this paper.

## Data availability statement

The data that support the findings of this study are available on request from the corresponding author from https://mynotebook.labarchives.com. RNAseq data is available at Table S2. Metabolomic data is available at Table S5.

## Authors’ Contributions

JF carried out the *in vitro* study using cell lines, contributed to the interpretation of the results and data analysis, took the lead in writing the manuscript with assistance from ZH, LT, and SKM. ZH was in charge of overall study direction and planning, supervised the project, edited the manuscripts, and contributed to the interpretation of the results. CR, ZH, OB, IC, and LT carried out the animal experiments. SKM edited the manuscripts. All authors provided critical feedback, helped shape the research, discussed the results, and contributed to the final manuscript.

## Acknowledgments

This project was supported by R03 funding, ID: 5R03AG068685-02, PI: Zhiqing Huang, by NIA/NIH, in 2021-2024. We sincerely express our gratitude to NIA/NIH. We are deeply grateful to all authors who contributed to the success of this research project. We would also like to thank our lab manager, Carole Grenier, who provided technical training and safety support for this research. We thank our undergrad students, Lukas Bleichner and Jocelyn Reyes, for checking the wellbeing of animals. We thank the undergrad students, Lila Teitle, Bella Mendieta, Hannah Lee, and Gaomong Lo, for the assistance of lab work. Finally, we would like to extend our heartfelt thanks to all the participants in our study, who generously shared their time, experiences, and insights with us.

## Notes

### Competing Interest Statement

The authors have declared no competing interest.

