## Supplemental information for "Regulation of Age-Related Lipid Metabolism in Ovarian Cancer"

**Supplementary information_ Quantitative Lipidomics Assay**

The whole procedure was carried out by Metware Biotechnology Inc. The fat tissues were homogenized by ball-mill in extraction solution (MTBE: Methanol = 3 : 1, V/V), vortexed, and incubated at 4°C for 10 min. After centrifugation at 12000 rpm for 10 min at 4°C, 200 µL of the upper phase was collected for complete solvent drying under 20°C. The residue was reconstituted using 200 uL of reconstitution solution (ACN: IPA = 1:1, V/V), followed by vortex and centrifugation. A 120 µL of the final supernatant was used for LC-MS analysis. The sample extracts were analyzed using an LC-ESI-MS/MS system (UPLC, Nexera LC-40, https://www.shimadzu.com; MS, Triple Quad 6500+, https://sciex.com/). The analytical conditions were as follows, UPLC: column, Thermo Accucore™C30 (2.6 µm, 2.1 mm×100 mm i.d.); solvent system, A: acetonitrile/water (60/40,V/V, 0.1% formic acid, 10 mmol/L ammonium formate), B: acetonitrile/isopropanol (10/90 VV/V, 0.1% formic acid, 10 mmol/L ammonium formate); gradient program, A/B (80:20, V/V) at 0 min, 70:30 V/V at 2.0 min, 40:60 V/V at 4 min, 15:85 V/V at 9 min, 10:90 V/V at 14 min, 5:95 V/V at 15.5 min, 5:95 V/V at 17.3 min, 80:20 V/V at 17.3 min, 80:20 V/V at 20 min; flow rate, 0.35 ml/min; temperature, 45℃; injection volume: 2 µL. The effluent was alternatively connected to an ESI-triple quadrupole-linear ion trap (QTRAP)-MS. LIT and triple quadrupole (QQQ) scans were acquired on a triple quadrupole-linear ion trap mass spectrometer (QTRAP), QTRAP® 6500+ LC-MS/MS System, equipped with an ESI Turbo Ion-Spray interface, operating in positive and negative ion mode and controlled by Analyst 1.6.3 software (Sciex). The ESI source operation parameters were as follows: ion source, turbo spray; source temperature 500 ℃; ion spray voltage (IS) 5500 V (Positive), -4500 V(Neagtive); Ion source gas 1 (GS1), gas 2 (GS2), curtain gas (CUR) was set at 45, 55, and 35 psi, respectively. Instrument tuning and mass calibration were performed with 10 and 100 μmol/L polypropylene glycol solutions in QQQ and LIT modes, respectively. QQQ scans were acquired as MRM experiments with collision gas (nitrogen) set to 5 psi. DP and CE for individual MRM transitions was done with further DP and CE optimization. A specific set of MRM transitions were monitored for each period according to the metabolites eluted within this period.

**Transcriptome Sequencing (RNA-Seq)**

The RNAseq was carried out using Roche Kapa Hyper RNA with Riboerase HMR library kit, and sequencing on Illumina NovaSeq X plus platform for 60M PE150 reads (30M reads each side). The quality of the sequencing output was first assessed by FastQC. Quality filtering and removal of residual adaptor sequences were conducted on read pairs using Trimmomatic while the reads longer than 36 bp were retained. STAR aligner was employed to align the reads with the reference of the rat genome Rnor 6.0 (GCA_000001895.4) and then Htseq-count package was utilized to generate gene read counts.

| Table 1 The IC50 alues of Drug in TSA-A&Y group | | | | |
|---|---|---|---|---|
| **Drug** | Young | Aged | Aged-Young | t test |
| **Olaparib** | 1941.389613 | 102.3773646 | -1839.012248 | 0.043451046 |
| **ZM447439** | 26.79720584 | 16.96608259 | -9.831123253 | 0.001030271 |
| **Palbociclib** | 530.8497063 | 45.39311237 | -485.4565939 | 0.031866839 |
| **Dactolisib** | 0.352917317 | 0.148255136 | -0.204662181 | 0.005533609 |
| **Pictilisib** | 5.310783666 | 3.259253621 | -2.051530044 | 0.000132558 |
| **PD0325901** | 52.91930707 | 3.270523065 | -49.648784 | 0.008417768 |
| **Rapamycin** | 0.260218466 | 0.086379998 | -0.173838468 | 0.014833941 |
| **Dabrafenib** | 548.1906076 | 61.80105827 | -486.3895493 | 0.002169162 |
| **AZD1208** | 272.4494186 | 159.8217316 | -112.627687 | 0.004595138 |
| **LCL161** | 996.2584154 | 120.5279726 | -875.7304428 | 0.013984863 |
| **Alpelisib** | 2778.378846 | 91.65077842 | -2686.728068 | 0.025068084 |
| **Taselisib** | 17.07113081 | 5.12641726 | -11.94471355 | 0.000139022 |
| **SCH772984** | 59.20484095 | 10.20236977 | -49.00247118 | 0.003172988 |
| **ERK_2440_1713** | 14.55180699 | 13.34899063 | -1.202816362 | 0.003643104 |
| **ERKERK_2440_1713** | 37.64593795 | 25.39086318 | -12.25507477 | 0.006048809 |
| **IRAK4** | 209.9553164 | 101.9879768 | -107.9673396 | 0.000402889 |
| **Carmustine** | 1972.822245 | 343.719145 | -1629.1031 | 0.006970552 |
| **Dactinomycin** | 0.264569657 | 0.057559959 | -0.207009698 | 0.003531957 |
| **Sinularin** | 167.3116606 | 24.98837894 | -142.3232816 | 0.005437135 |
| **Ulixertinib** | 4181.530001 | 96.23721486 | -4085.292786 | 0.03454911 |
| **AZD3759** | 17.48824429 | 12.20536131 | -5.282882976 | 0.001614703 |
| **Cediranib** | 8.023990593 | 13.97362786 | 5.949637272 | 0.022510932 |
| **GSK2578215A** | 504.4844424 | 96.24063513 | -408.2438073 | 0.024572418 |
| **P22077** | 128.2021556 | 68.86954563 | -59.33261 | 0.002366603 |
| **Foretinib** | 37.8235241 | 3.092357616 | -34.73116648 | 0.005335262 |
| **AZ6102** | 19.84322806 | 8.496260154 | -11.34696791 | 0.000311793 |
| **GSK591** | 4249.041367 | 204.1732633 | -4044.868104 | 0.041619162 |

**Table S1. TaqMan probes for RT-qPCR** (Manufacturer: ThermoFisher)

| **Gene ID** | **Cat #** |
|---|---|
| S100a8 | Rn00587579_g1 |
| S100a9 | Rn00585879_m1 |
| Il1rl1 | Rn01640664_m1 |
| Lcn2 | Rn00590612_m1 |
| C3 | Rn00566466_m1 |
| Hba-a1 | Rn01789798_s1 |
| Fcna | Rn00580247_m1 |
| Pnpla3 | Rn01502361_m1 |
| GAPDH | Rn01775763_g1 |
